# Assessment of chimeric antigen receptor T (CAR-T) cytotoxicity by droplet microfluidics *in vitro*

**DOI:** 10.1101/2021.12.28.474351

**Authors:** Kuan Un Wong, Jingxuan Shi, Peng Li, Haitao Wang, Yanwei Jia, Chuxia Deng, Lianmei Jang, Ada Hang-Heng Wong

## Abstract

Chimeric antigen receptor T (CAR-T) cells are cytotoxic T cells engineered to specifically kill cancer cells expressing specific target receptor(s). Prior CAR-T efficacy tests include CAR expression analysis by qPCR or ELISA, *in vitro* measurement of interferon-γ (IFNγ) or interleukin-2 (IL-2), and xenograft models. However, the *in vitro* measurements did not reflect CAR-T cytotoxicity, whereas xenograft models are low throughput and costly. Here we presented a robust *in vitro* droplet microfluidic assay for CAR-T cytotoxicity assessment. This method not only enabled assessment of CAR-T cytotoxic activity under different fluid viscosity conditions, but also facilitated measurement of CAR-T expansion and dissection of mechanism of action via phenotype analysis *in vitro*. Furthermore, our data suggested that label-free cytotoxicity analysis is feasible by acquiring data before and after treatment. Hence, this study presented a novel *in vitro* method for assessment of cellular cytotoxicity that could potentially be applied to any cell-kill-cell experiment with varying solvent composition.

## Introduction

Chimeric antigen receptor T (CAR-T) cells have been approved by the US Food and Drug Administration (FDA) for application in cancer immunotherapy. Prior CAR-T efficacy tests are summarized in Table 1.

**Table 1.**
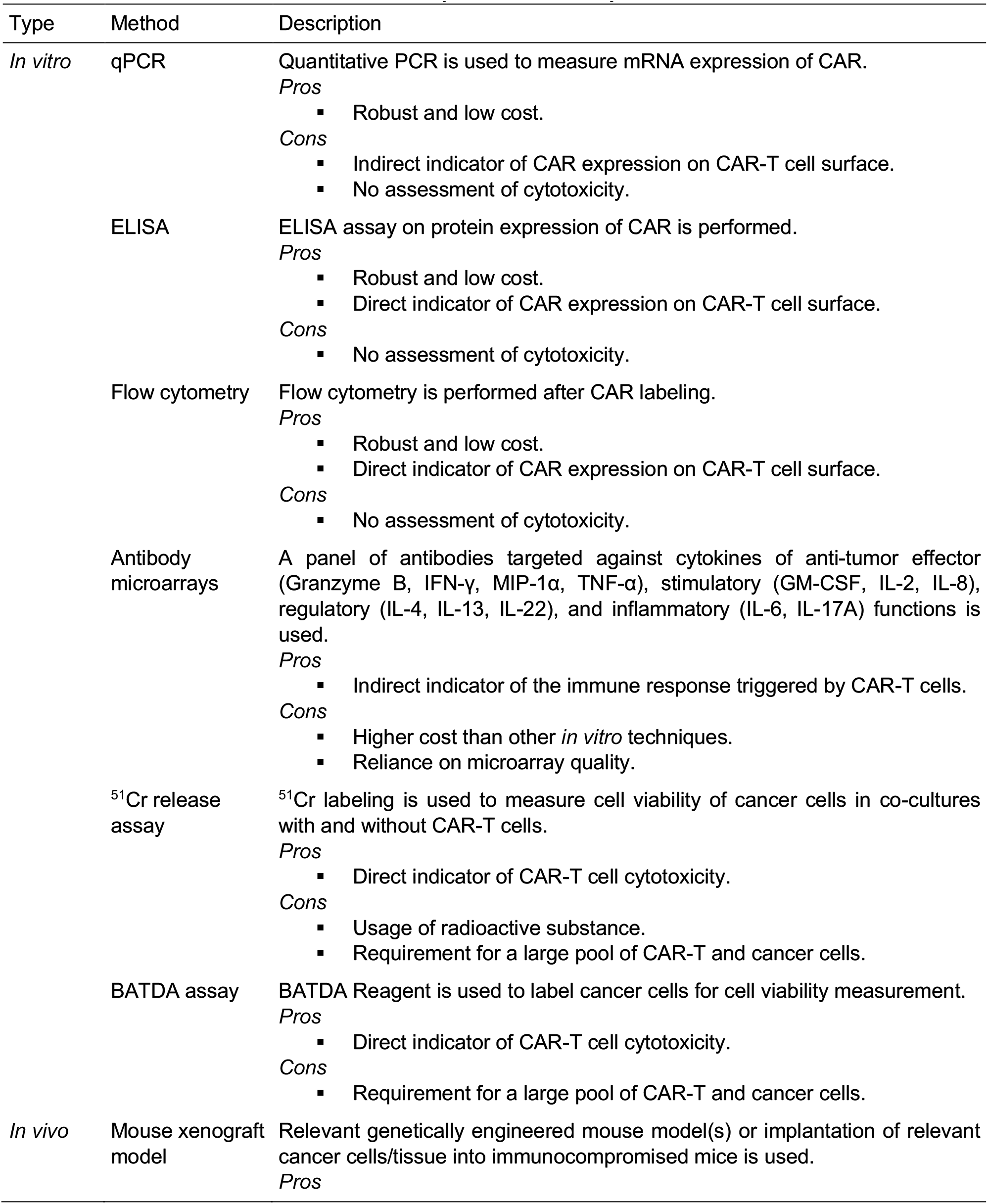
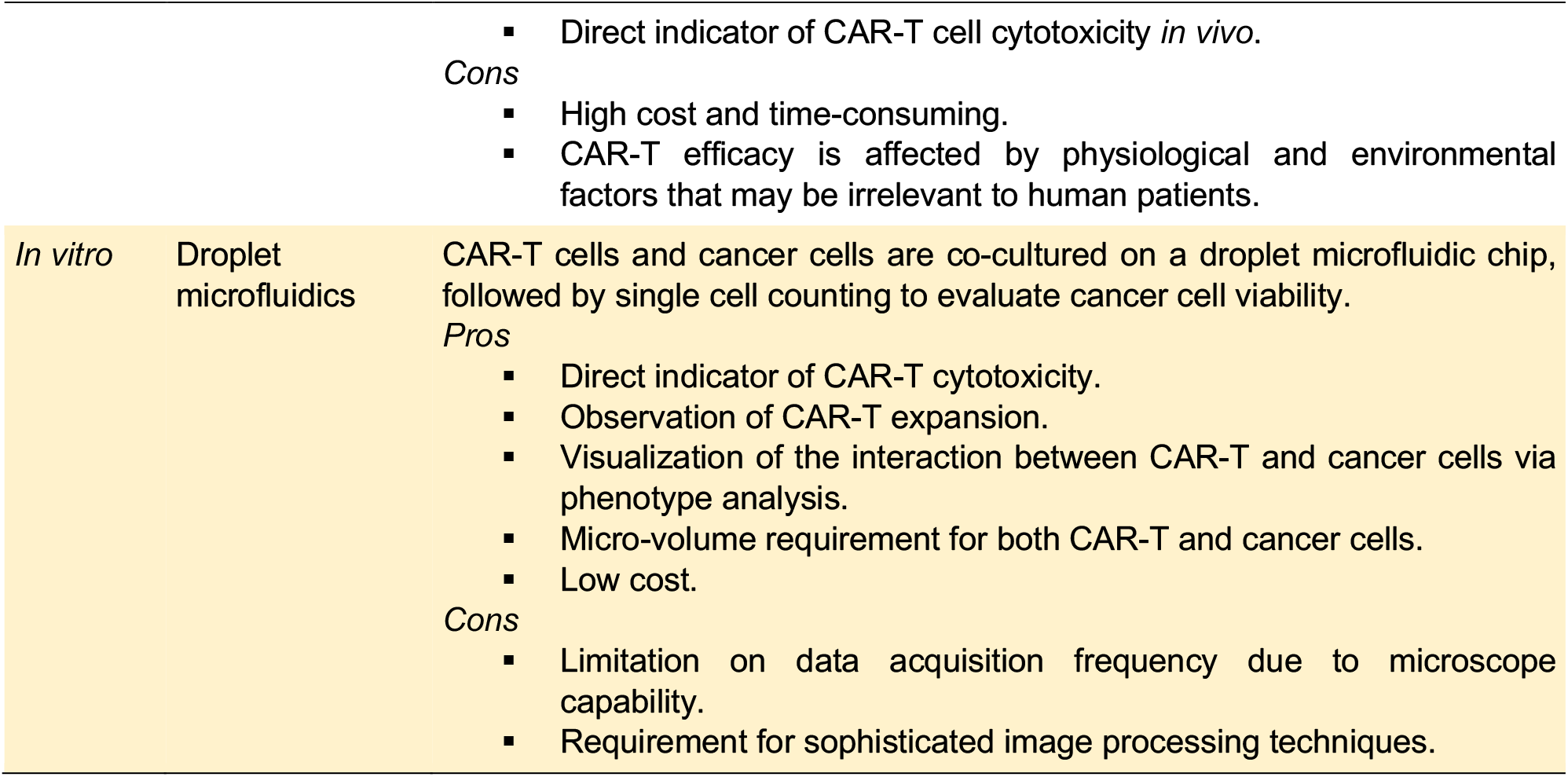
Summary of CAR-T efficacy tests ^1,2^.

The major drawback of current CAR-T cytotoxicity assessment methods is that a large pool of both the CAR-T and cancer cells is required, especially for the xenograft models.

Recently, novel microfluidic technologies enable cytotoxicity assessment using different cytotoxic cells as model ^3^. Among these microfluidic assays, the T-or Y-channels are very robust droplet generation systems, but the low cell occupancy of the majority of these systems impedes real-life applications. First, >99% of droplets are empty when these systems are tailored to obtain one cell per droplet unless very specific conditions can be met ^4^. Second, it is difficult to precisely quantify droplets maintained in large, open chambers ^5^. Third, confining droplets into microwells ^6^ may aid time-lapse imaging and effective clone selection from a pool, but the propensity of empty droplets will greatly increase workload if all cells of a tumor need to be analyzed to address heterogeneity. In contrast, our microfluidic chip ^7^ enables 100% microwell occupancy and 100% cell-containing droplets when used appropriately. Additionally, the large droplets allow imaging of cell morphology and movement at multiple time points, which cannot be achieved by cell traps ^8^. Hence, our chip assay minimizes empty droplet incidences and provides cell morphology information that other systems could not achieve.

Alternatively, as opposed to microwell microfluidic systems, our chip enables flexible sample replicates (i.e. the number of loaded microwells) without the need to fabricate new chips. This flexibility can be partially achieved by a microfabricated 96-well plate ^9^, if there is sufficient sample to fill at least one well. Additionally, these plates are relatively costly unless the full plate is used at the same time, whereas our chip is designed for one sample per chip to ensure sterile condition and avoid cross-contamination. Alternatively, in open microwell systems, adjusting cell concentration to adjust sample size will not only affect cell distribution per microwell, but also microwell occupancy rate, the latter of which is unaffected in our system unless one cell per drop is desired. Micro-patterning potentially increases microwell occupancy in microwell systems, but it complicates chip fabrication and increases cost. Therefore, our chip assay provides loading flexibility while retaining screening efficacy.

On the other hand, more sophisticated systems have been built to investigate the invasiveness of therapeutic cells through biogels ^10–12^. However, quantification in these assays will be compromised. Furthermore, many systems were developed to address the issue of cytokine production ^13–15^, but most of these systems are either delinked from quantitative cytotoxicity assessment or impractical for multi-timepoint analysis. In conclusion, each assay has its pros and cons. Our chip design provides genuine advantage for rapid cell-based cytotoxicity screens applied to cells that are adaptable to suspension culture.

In this study, we exploited our previously developed droplet microfluidic chip ^7^ to establish a two-cell type model for screening CAR-T cells against cancer cells. The basic chip design and data acquisition procedures remained the same, but cell type identification became challenging. Eventually, we tested the application using a CD19^+^ Nalm6 cancer cell line stably expressing green fluorescent protein (GFP) and luciferase ^18^ to setup our model. We analyzed two batches of CAR-T cells *in vitro* using luciferase assay and our chip assay, where luciferase activity measurements were calibrated by a standard curve plotted by gradient Nalm6 cell numbers against blank control (Figure S1). The two CAR-T batches demonstrated similar cytotoxic activity by luciferase activity, but significantly different cytotoxicity on chip. Therefore, we performed in-depth analysis of the chip assay to exemplify the capability of this method.

## Materials and Methods

### Preparation of chimeric antigen receptor T (CAR-T) cells

Usage of primary human T cells in this study was approved by the Institutional Review Board of Guangzhou Institute of Biomedicine and Health, Chinese Academy of Sciences, under ethics approval no. GIBH-IRB07-2018024. Peripheral mononuclear cells (PBMCs) were separated in Lymphoprep™ (STEMCELL™ Technologies) by density gradient centrifugation. Primary human T cells were isolated from PBMCs by negative selection using the Pan T Isolation Kit (Miltenyi Biotec). Batch 1 T cells were cultured in IMDM Medium containing Icove’s Modified Dulbecco’s Medium (IMDM) (Life Technologies) supplemented with 10% (v/v) fetal bovine serum (Life Technologies) and 100 U/mL Penicillin-Streptomycin (Life Technologies), and stimulated with TransAct™ (Miltenyi Biotec). Batch 2 T cells were cultured in R10 Medium containing RPMI 1640 (Life Technologies) supplemented with 10% fetal bovine serum (Life Technologies), 100 U/mL Penicillin-Streptomycin (Life Technologies), 2 mM L-glutamine (Life Technologies), 25 mM HEPES pH7.5 (Life Technologies) and 100 μg/mL Streptavidin (Sigma-Aldrich), and stimulated with particles coated with anti-CD3/anti-CD28 antibodies (Miltenyi Biotec) at a cell-to-bead ratio of 1:2. Approximately 48 h after activation, T cells were transfected with supernatant containing lentiviral vector expressing CAR. After transduction for 12 h, all transduced T cells were sorted by fluorescence activated cell sorting (FACS) using anti-EGFR antibody (BioLegend). Purified CAR-T cells were cultured or assayed with Activation Medium containing IMDM Medium or R10 Medium supplemented with 300 U/mL IL-2 (clinical grade). The CAR-T cells were fed every 2 days with fresh media for no more than 5 d, and frozen in liquid nitrogen until use following standard freezing procedures.

### Preparation of Nalm6 cancer cells stably expressing green fluorescent protein (GFP)

Nalm6 cells (ATCC^®^) were transduced with a lentiviral vector co-expressing GFP and luciferase. Nalm6 cells were cultured in Culture Medium containing RPMI 1640 (Life Technologies) supplemented with 10% (v/v) fetal bovine serum (Life Technologies), 100 U/mL Penicillin-Streptomycin (Life Technologies), 2 mM L-glutamine (Life Technologies), 25 mM HEPES pH7.5 (Life Technologies) and 50 μM β-mercaptoethanol (Sigma-Aldrich).

### Luciferase assay

1.0 × 10^5^ Nalm6 cells were incubated with CAR-T cells in Activation Medium at the indicated ratios in triplicate wells of a 96-well U-bottomed plate for 18 h. Luminescence was obtained after applying the Dual-Luciferase Assay Kit (Promega) in a 96-well opaque plate following manufacturer’s protocol and measured on Perkin Elmer Victor X3 Plate Reader. Background luminescence was negligible (signal : noise (S/N) ratio <0.01). Subsequently, cell viability was calculated by the following formula:

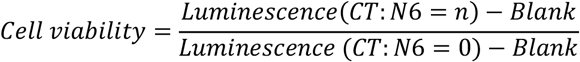

For comparison to on-chip results, a standard curve of serially diluted Nalm6 cells from 2.0 × 10^5^ cells/well to 3.1 × 10^3^ cells/well was plotted against luminescence. Linear regression was performed using Microsoft Excel 2013 for calculation of absolute cell numbers from relative luminescence.

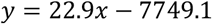

where *y* denoted luminescence and *x* denoted Nalm6 cell number.

Blank controls contained Culture Medium but no cells.

### On-chip CAR-T cytotoxicity assay

CAR-T and Nalm6 cells were mixed at a final concentration of 5.0 × 10^6^ and 1.0 × 10^6^ cells/mL in Activation Medium, respectively. 1% (w/v) methyl cellulose (Sigma-Aldrich) was added to high viscosity medium (HiVis), whereas the same volume of ultrapure water was added to low viscosity medium (LoVis). 2 μM ethidium homodimer 1 (Life Technologies) was added to stain dead cells if indicated. Chip loading was performed as previously described ^7^. Chips were incubated in a 37°C humidified incubator supplemented with 5% CO_2_. All wells on chip were imaged under 10× objective in the channels of brightfield, GFP and RFP on Life Technologies EVOS FL Imaging System at 0 h and 24 h post-treatment respectively.

### Bioimaging assay

One polydimethylsiloxane (PDMS) well was fabricated at the center of a 35 mm cell culture dish by mixing 1:10 (w/w) PDMS catalyst : substrate (Silmore) followed by incubation at 65°C overnight. Fabricated dishes were kept in a sealed box at room temperature until use.

1.0 × 10^6^ CAR-T cells and 2.0 × 10^5^ Nalm6 cells were loaded into the PDMS well of a fabricated dish, followed by addition of 1.5 mL Activation Medium to fill the dish. Next, the dish was calibrated in the 37°C humidified chamber supplemented with 5% CO_2_ of Nikon Biostation IM-Q Imaging System for 30 min. Time-lapse imaging at 10× or 20× magnification was performed at 2-10 min intervals for no more than 24 h. Raw image sequences were exported as TIFF.

### Data analysis and visualization

All images were initially imported as image stacks into ImageJ v1.52e, followed by brightness and contrast adjustment, cropping, trimming, and image stack reduction to obtain the final image sequence or montage as depicted. Video clips were edited by CyberLink PowerDirector 17 to include annotations, followed by export to MPEG4. All graphs and plots were drawn by GraphPad Prism v5.1 or using *ggplot* in R v3.3.2. Statistical analysis was performed as indicated in Data S1 or in corresponding figure legend(s). Figures were prepared by assembling images, graphs and plots using Adobe^®^ Illustrator^®^ CS6 v16.0.0.

## Results

We established our model using anti-CD19 CAR-T cells (Figure 1a). In our assay, CAR-T and Nalm6 cells were initially mixed in Activation Medium at a cell number ratio of 5:1, followed by chip loading and imaging before and after incubation for 24 h (Figure 1b). Addition of 1% methyl cellulose, which increased fluid viscosity and facilitated adherent cell suspension on chip ^7^, was conducted to observe cell behavior under different fluid viscosity conditions. Eventually, the cells were identified, classified and counted before and after incubation, followed by calculation of cell viability and CAR-T:Nalm6 (CT:N6) ratio in each droplet (Figure 1c).

**Figure 1.**
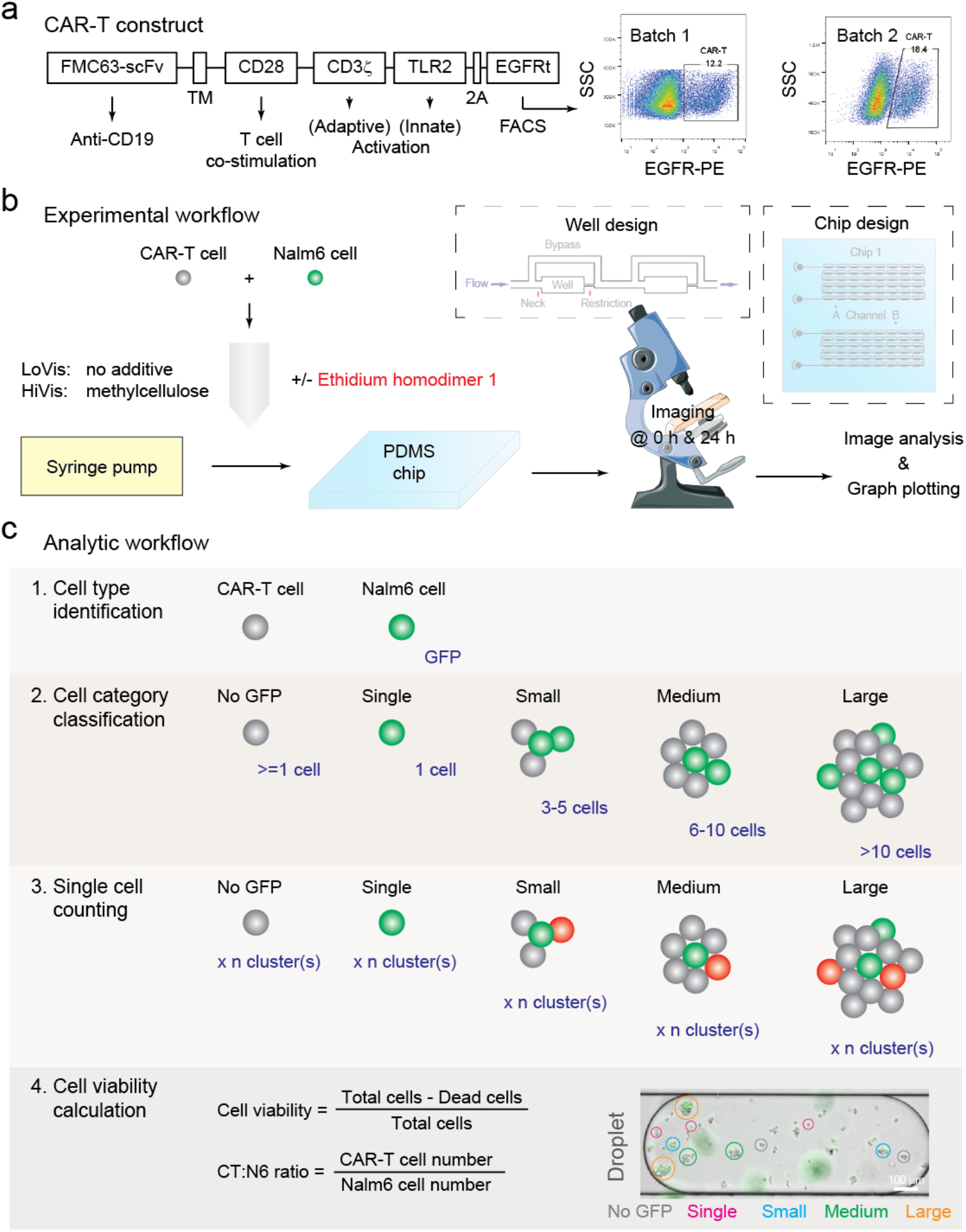
Paradigm of this study. (**a**) CAR-T construct design and FACS characterization. CAR was constructed with the CD19-targeting FMC63-scFv domain (FMC63-sFv), the CD28 transmembrane domain (TM), the CD28 endodomain (CD28), CD3ϋ signaling domain (CD3ϋ), the Toll/interleukin-1 receptor domain of Toll-like receptor 2 (TLR2), the 2A self-cleaving peptide (2A) and the extracellular and transmembrane domains of truncated human epidermal growth factor receptor (EGFRt). CAR-T cells were purified 12 h post-transduction using anti-human EGFR antibody and side scattering (SSC); the percentage of CAR-T cells in the population of all transduced T cells was indicated. (**b**) Experimental workflow of this study. CAR-T and Nalm6 cancer cells were mixed and loaded on chip, followed by 24 h incubation and imaging. The design of the chip and its wells was shown in the insets. (**c**) Analytic workflow of this study. Nalm6 cells were initially identified by GFP fluorescence, followed by cell category classification, cell counting and viability calculation. Cell category classification was arbitrary as shown in the inset. Dead cells were indicated by the red fluorescent dye ethidium homodimer 1 (EH1); for EH1-free assays, cell viability was calculated by the difference of the number of Nalm6 cells before and after treatment.

### Validation of chip performance

Because cell-cell interaction has never been studied on our chip, we initially verified the performance of our droplet microfluidic chip for the quantitative measurement of CAR-T cytotoxicity. Firstly, we validated the cell number distribution frequency in each droplet. In consistence to our one-cell type model ^7^, the number of each cell type in each droplet displayed Poisson distribution in low viscosity medium without additive and in high viscosity medium containing methyl cellulose, respectively (Figures 2a & 2b). Notably, no well contained 0 CAR-T cells under both fluid conditions, implying that no Nalm6 cell had nil probability to interact with CAR-T cells. Secondly, the total number of the two cell types in each droplet displayed Poisson distribution under both fluid conditions (Figure 2c). Thirdly, the CT:N6 ratio in each droplet displayed Gaussian distribution under both fluid conditions (Figure 2d). Cell number and ratio distribution patterns was conserved in Batch 2 CAR-T cells (Figure S2). Collectively, these data confirmed the statistical patterns of cell distribution in the two-cell type model.

**Figure 2.**
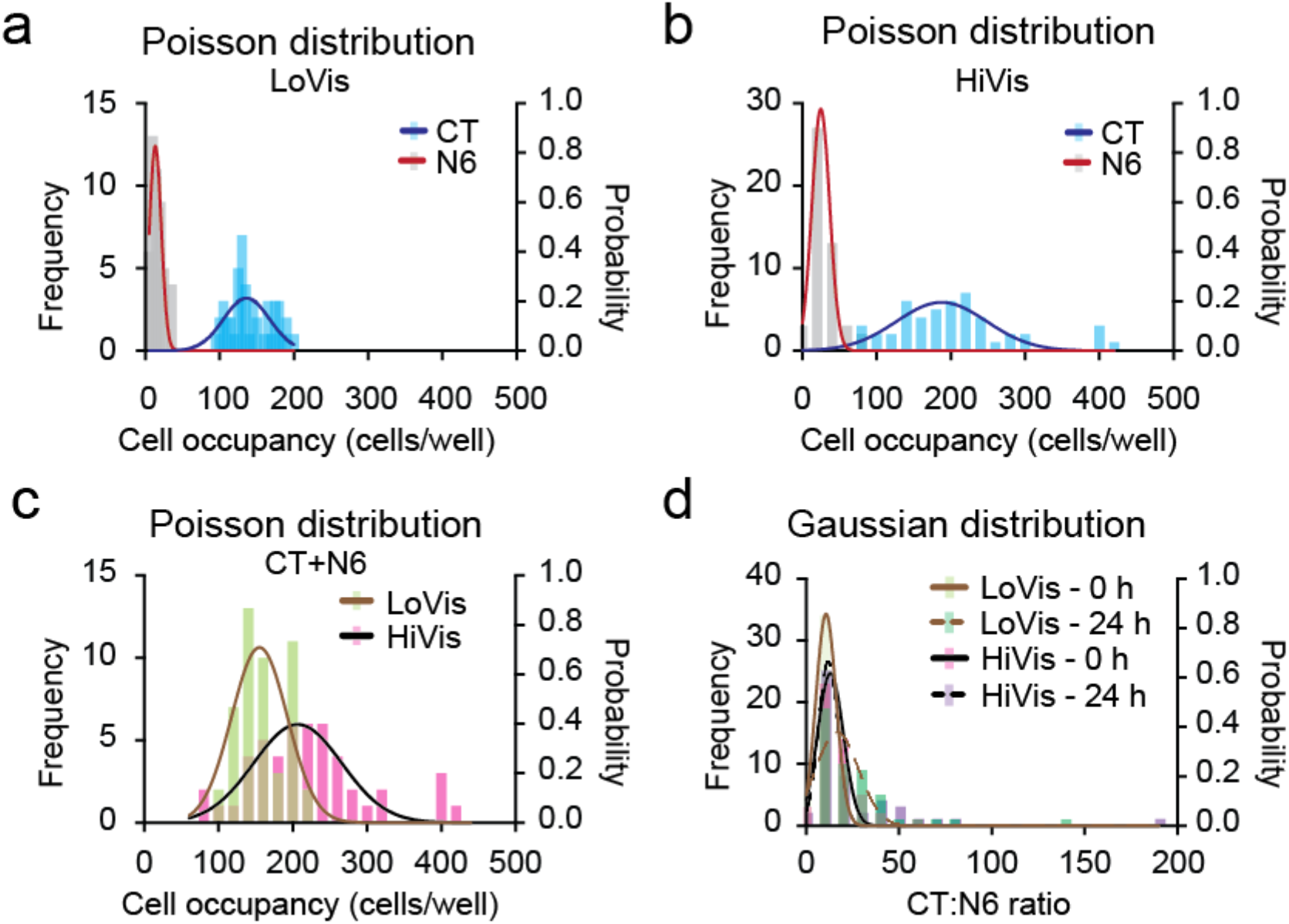
Cell occupancy analysis of Batch 1 CAR-T cells. (**a-c**) Poisson distribution of Batch 1 CAR-T and Nalm6 cells on chip. Frequency distribution analysis of CAR-T (CT) and Nalm6 (N6) cells in low viscosity medium (LoVis) and high viscosity medium (HiVis) on chip at 0 h was performed. Distribution frequency and probability was plotted against cell occupancy in all wells on chip; each column represented one data grid automatically classified by GraphPad Prism; the lines indicated Poisson distribution curves generated by GraphPad Prism. (**d**) Gaussian distribution of CT:N6 ratio on chip. Frequency distribution analysis of CT:N6 ratio under both fluid conditions before and after treatment was performed. Distribution frequency and probability was plotted against CT:N6 ratio in all wells on chip; each column represented one data grid automatically classified by GraphPad Prism; the lines indicated Gaussian distribution curves generated by GraphPad Prism.

### Analysis of CAR-T cytotoxicity on chip

After confirming chip performance, we started to analyze CAR-T cytotoxicity on chip.

Firstly, we analyzed the cytotoxicity of Batch 1 CAR-T cells against Nalm6 cells. Taking EH1-labelled cells into account, results showed that the overall droplet cell viability of Nalm6 cells was 53.6% and 100.7% in low viscosity medium and high viscosity medium respectively (Figure 3a). To exploit our assay to its full potential, we dissected the cell killing paradigm under each fluid condition after classifying cells into arbitrary cell categories (Figure 1c). Analysis of Nalm6 occupancy depicted that Nalm6 cells were evenly distributed in all cell categories without significant difference under both fluid conditions (Table S1). However, significant CAR-T cytotoxicity occurred in single (p = 9.5e-11), small (p = 5.0e-6) and medium clusters (p = 1.5e-3) but not in large clusters (p = 4.7e-1) in low viscosity medium (Figure 3b), whereas CAR-T cytotoxicity was only significant in single (p = 1.2e-5) and small clusters (p = 6.7e-7) in high viscosity medium (Figure 3c), suggesting that CAR-T cytotoxicity diminished in larger clusters as compared to smaller ones.

**Figure 3.**
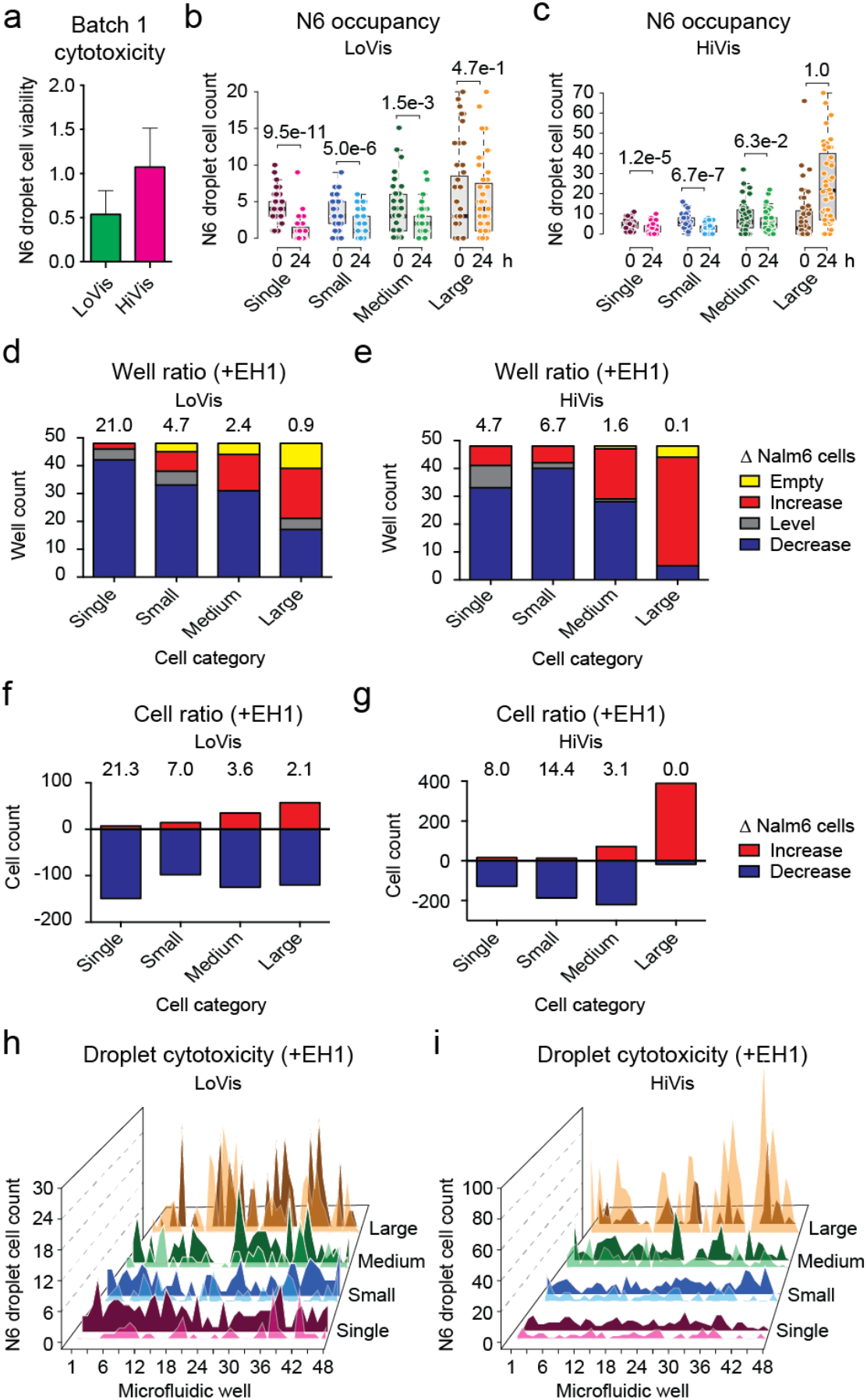
Cytotoxicity assessment of Batch 1 CAR-T cells. (**a**) Overall cytotoxicity of Batch 1 CAR-T cells under different fluid conditions. Mean Nalm6 droplet cell viability was plotted against low viscosity medium (LoVis) and high viscosity medium (HiVis) respectively; the error bars denoted standard deviation of all replicate wells. (**b**,**c**) Analysis of Nalm6 occupancy before and after treatment on chip. Analysis of Nalm6 occupancy in different cell categories at 0 h and 24 h post-treatment on chip was performed. Nalm6 droplet cell count was plotted against cell category and treatment time; each dot represented one well; the thick black line, grey box and dotted error bars of the box plot indicated median Nalm6 droplet cell count, interquartile range between -1α and 1α, and range of Nalm6 droplet cell count of all wells, respectively; the numbers indicated p-values of 2-tailed *t* test. (**d**,**e**) Well ratio analysis of CAR-T cytotoxicity under different fluid conditions. Well ratio was plotted against cell categories; each column indicated the number of wells containing no, increasing, same or decreasing number of Nalm6 cells; the numbers indicated the well ratio of the specified dataset. (**f**,**g**) Cell ratio analysis of CAR-T cytotoxicity under different fluid conditions. Cell ratio was plotted against cell categories; each column indicated the number of Nalm6 cells that increased or decreased in sum; the numbers indicated the cell ratio of the specified dataset. (**h**,**i**) Hill plot of Nalm6 occupancy along all wells on chip. Nalm6 droplet cell count was plotted against consecutive wells on chip; each peak indicated the number of Nalm6 cells in the specified droplet; dark-colored shade indicated the cell number before treatment, whereas light-colored shade indicated the cell number after treatment.

To further dissect the mechanism, we summarized CAR-T cytotoxicity by the well ratio and the cell ratio that are correlated to Nalm6 cell number variance in droplets. The well ratio, expressed as the number of wells with decreasing Nalm6 cells to that of increasing Nalm6 cells, suggested of CAR-T cytotoxicity in a population of droplets. Alternatively, the cell ratio, expressed as the number of decreased Nalm6 cells divided by the number of increased Nalm6 cells, suggested of the cytotoxic activity of CAR-T cells as a population. Analysis of CAR-T cytotoxicity in different cell categories showed that single cells (well ratio = 21.0, cell ratio = 21.3) demonstrated exquisite CAR-T cytotoxicity in low viscosity medium (Figures 3d & 3f), whereas both single cells (well ratio = 4.7, cell ratio = 8.0) and small clusters (well ratio = 6.7, cell ratio = 14.4) depicted good CAR-T cytotoxicity in high viscosity medium (Figures 3e & 3g). Conversely, large clusters showed poor CAR-T cytotoxicity under both fluid conditions (Figures 3d - 3g).

Next, we tested if Nalm6 occupancy or CAR-T cytotoxicity could be correlated to well position on chip, which is critical to multi-drug loading ^7^. First, no apparent distribution of Nalm6 occupancy was observed along the wells on chip under both fluid conditions (Figures 3h & 3i). Second, two-way ANOVA analysis of Nalm6 occupancy in different cell categories against well position showed that there was insignificant difference between Nalm6 occupancy and well position in all cell categories in low viscosity medium (p = 0.291), whereas minor correlation arisen from extreme Nalm6 cell numbers as demonstrated by Tukey’s honestly significant difference (HSD) test was observed in high viscosity medium (p = 0.047) (Table S2). Therefore, we deduced that Nalm6 occupancy in droplets and in clusters are both uncorrelated to well position. Third, two-way ANOVA analysis of the difference of Nalm6 occupancy between 0 h and 24 h post-treatment against well position showed that there was no significant correlation between CAR-T cytotoxicity and well position in low viscosity medium (p = 0.941) and high viscosity medium (p = 0.964) respectively (Table S3). In contrast, variant CAR-T cytotoxicity among different cell categories led to significant difference of Nalm6 occupancy after 24 h treatment in high viscosity medium (p < 0.0001) but not in low viscosity medium (p = 0.186) (Table S3). This observation was consistent to the disparity between the ratiometric parameters of the two fluid conditions that CAR-T cells as a population were still actively killing Nalm6 cells in large clusters in low viscosity medium, whereas Nalm6 cell proliferation dominated over CAR-T cytotoxicity in large clusters in high viscosity medium (Figures 3d - 3g). Collectively, these data suggested of the lack of global modulation factor(s) that coherently affect CAR-T cytotoxicity within one droplet. This suggested that the droplet merely defines the spatial constraint of cell-cell interaction, whereas the cell clusters in each droplet provide micro-reaction centers for CAR-T cytotoxic activity.

Hence, this method enables robust assessment CAR-T cytotoxicity and dissection of mechanism of action via phenotype analysis *in vitro*.

### Comparison between analysis with and without EH1 labelling

Ethidium homodimer 1 (EH1) was used as a dead cell indicator dye in our previous study ^7^. Although EH1 exhibited unapparent cytotoxicity in primary tumor dissociated cells in the investigated time frame ^7^, we tested if analysis without consideration of EH1 labelling influenced data interpretation to explore the feasibility of label-free cytotoxicity assessment.

In comparison to analysis with EH1 labelling, Nalm6 droplet cell viability was slightly higher without consideration of EH1 labelling, measured as 62.4% and 148.5% for the two fluid conditions, respectively (Figure 4a). This phenomenon is reasonable because EH1 indicates cells with compromised membrane integrity prior to death, so these cells are counted as viable if EH1 labelling is neglected. Concordantly, comparing Nalm6 droplet cell viability with or without consideration of EH1 labelling demonstrated that the difference was statistically insignificant in low viscosity medium (p = 1.4e-20) and high viscosity medium (p = 4.1e-19) (Table S4). Similarly, comparison between Nalm6 occupancy in different cell categories between the two analytic methods showed that there was insignificant difference at 0 h and 24 h under both fluid conditions (Table S5). In concordance, the well ratio and cell ratio followed similar patterns but lower absolute values in analysis without EH1 as compared to analysis with EH1 (Figure 4b). Hence, we concluded that EH1 labelling is dispensable for assessment of CAR-T cytotoxicity using our droplet microfluidic chip. In multi-drug assessment, we rationalize that drug ranking would remain the same on identical criteria. This conclusion thus implies that label-free assessment solely by cell counting before and after treatment is feasible.

**Figure 4.**
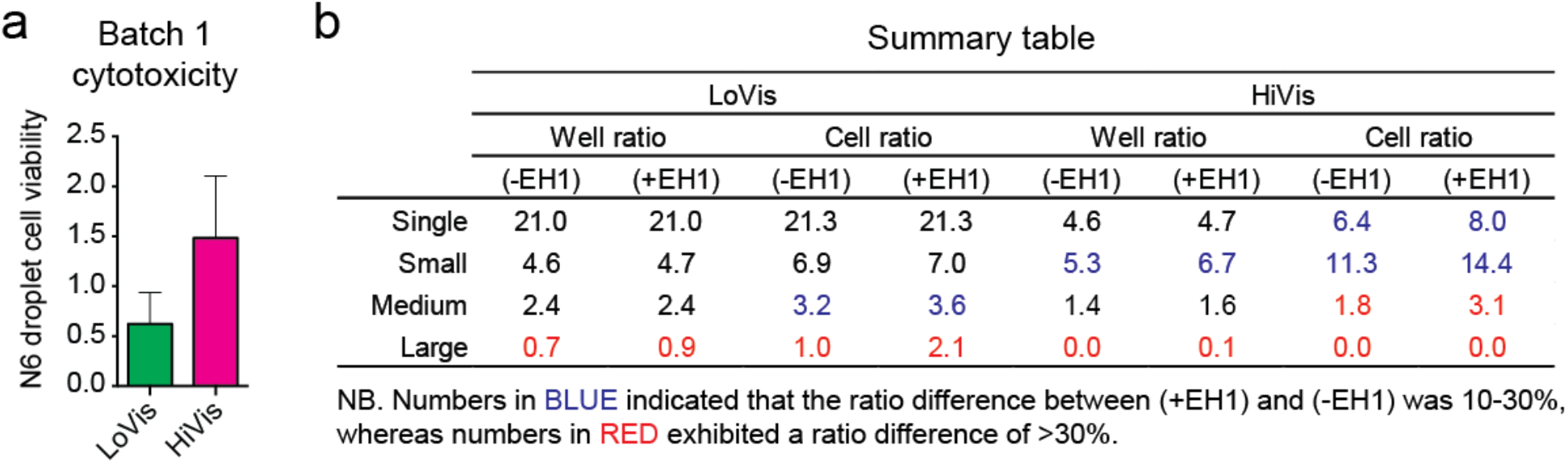
Cytotoxicity assessment of Batch 1 CAR-T cells without consideration of EH1 labelling. (**a**) Overall cytotoxicity of Batch 1 CAR-T cells under different fluid conditions. Mean Nalm6 droplet cell viability was plotted against low viscosity medium (LoVis) and high viscosity medium (HiVis) respectively; the error bars denoted standard deviation of the mean. (**b**) Comparison of well ratio and cell ratio with and without consideration of EH1 labelling under different fluid conditions. The well ratio and cell ratio of different cell categories obtained with and without consideration of EH1 labelling in low viscosity medium (LoVis) and high viscosity medium (HiVis) was summarized.

### Application on quality control of different batches of CAR-T cells

Given that label-free assessment is feasible, we tested the cytotoxicity of Batch 2 CAR-T cells without EH1 addition. First, although Batch 2 CAR-T cells were reasonably active in luciferase assay as compared to Batch 1 CAR-T cells (Figure S1), they showed significantly lower cytotoxicity (Figure 5a) as compared to Batch 1 CAR-T cells on chip (Figure 3a). Similarly, CAR-T cytotoxicity became insignificant in all cell categories under both fluid conditions (Figures 5b & 5c), reminiscent of the poor cytotoxic activity of Batch 2 CAR-T cells. Second, Batch 2 CAR-T cells demonstrated similar cytotoxic patterns in different cell categories (Figures 5d - 5g) as compared to Batch 1 CAR-T cells (Figures 3d - 3g), suggesting that the mechanism of action is unaffected by cytotoxic activity. Third, Nalm6 occupancy was also evenly distributed along all wells on chip (Figures 5h & 5i). Taken together, the well ratio and cell ratio reflected the variant cytotoxic activity and conserved mechanism of action between Batch 1 and Batch 2 CAR-T cells in this study. Hence, we interpreted the discrepancy of cytotoxic activity of Batch 2 CAR-T cells as a result of artefactual potency in luciferase assay due to exponential decay of bioluminescence. Our results suggested that our microfluidic chip assay can be applied to quality control between different batches of CAR-T cells.

**Figure 5.**
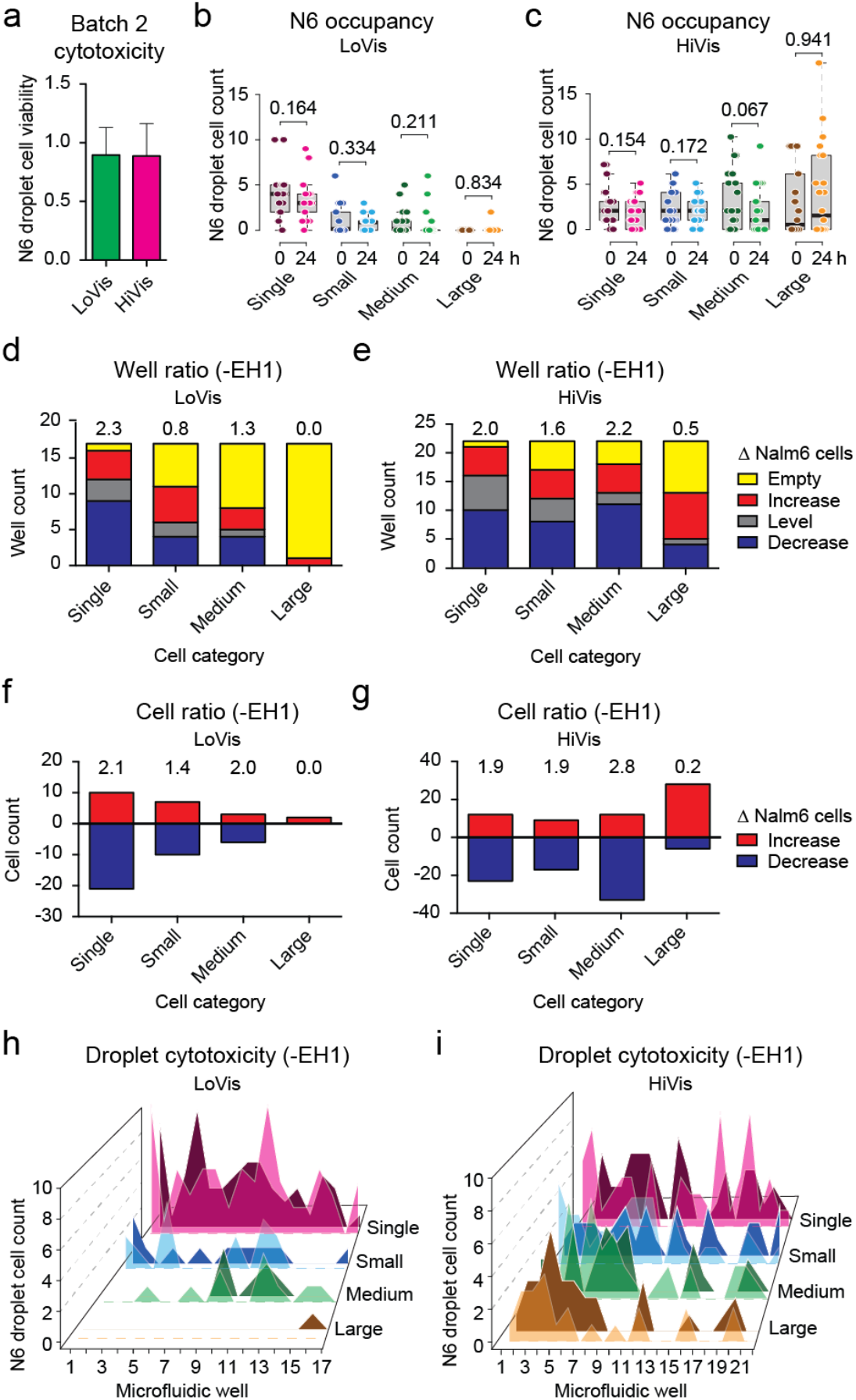
Cytotoxicity assessment of Batch 2 CAR-T cells. (**a**) Overall cytotoxicity of Batch 2 CAR-T cells under different fluid conditions. Mean Nalm6 droplet cell viability was plotted against low viscosity medium (LoVis) and high viscosity medium (HiVis) respectively; the error bars denoted standard deviation of all replicate wells. (**b**,**c**) Analysis of Nalm6 occupancy before and after treatment on chip. Analysis of Nalm6 occupancy in different cell categories before and after treatment on chip was performed. Nalm6 droplet cell count was plotted against cell category and treatment time; each dot represented one well; the thick black line, grey box and dotted error bars of the box plot indicated median Nalm6 droplet cell count, interquartile range between -1σ and 1σ, and range of all wells, respectively; the numbers indicated p-values of 2-tailed *t* test. (**d**,**e**) Well ratio analysis of CAR-T cytotoxicity under different fluid conditions. Well ratio was plotted against cell categories; each column indicated the number of wells containing no, increasing, same or decreasing number of Nalm6 cells; the numbers indicated the well ratio of the specified dataset. (**f**,**g**) Cell ratio analysis of CAR-T cytotoxicity under different fluid conditions. Cell ratio was plotted against cell categories; each column indicated the number of Nalm6 cells that increased or decreased in sum; the numbers indicated the cell ratio of the specified dataset. (**h**,**i**) Hill plot of Nalm6 occupancy along all wells on chip. Nalm6 droplet cell count was plotted against consecutive wells on chip; each peak indicated the number of Nalm6 cells in the specified droplet; dark-colored shade indicated the cell number before treatment, whereas light-colored shade indicated the cell number after treatment.

### Assessment of CAR-T cell expansion

CAR-T cell expansion is an important feature of efficient CAR-T cytotoxicity ^19,20^, but cannot be achieved in one assay using current protocols ^1,2^. For instance, Batch 1 CAR-T cells expanded in low viscosity medium (mean = 24.1, median = 18.5) and diminished in high viscosity medium (mean = -73.6, median = -73.5) with significant difference between fluid conditions (p < 0.0001) (Table S6). However, Batch 2 CAR-T cells diminished in low viscosity medium (mean = -3.2, median = -2.0) and high viscosity medium (mean = -6.8, median = -4.5) without significant difference between fluid conditions (p = 0.400) (Table S7). Hence, expansion of Batch 1 CAR-T cells was significant in low viscosity medium (p = 2.24e-6) but not in high viscosity medium (p = 1.00) (Figure 6a), in accordance to the measurement of Nalm6 cell reduction that reflected CAR-T cytotoxicity under these fluid conditions (Figure 3a). Consistently, Batch 2 CAR-T cell expansion (Figure 6b) was also reflective of its cytotoxic activity (Figure 5a). Therefore, our method not only enabled assessment of CAR-T cytotoxicity by Nalm6 occupancy, but also illustrated CAR-T cell expansion as supporting evidence. Although *in vitro* CAR-T cell expansion could not guarantee expansion *in vivo*, this method discriminates CAR-T cytotoxicity into the facets of cancer cell reduction and CAR-T cell expansion, which potentially facilitates in-depth investigation of CAR-T cytotoxic activity by the reductionist approach.

**Figure 6.**
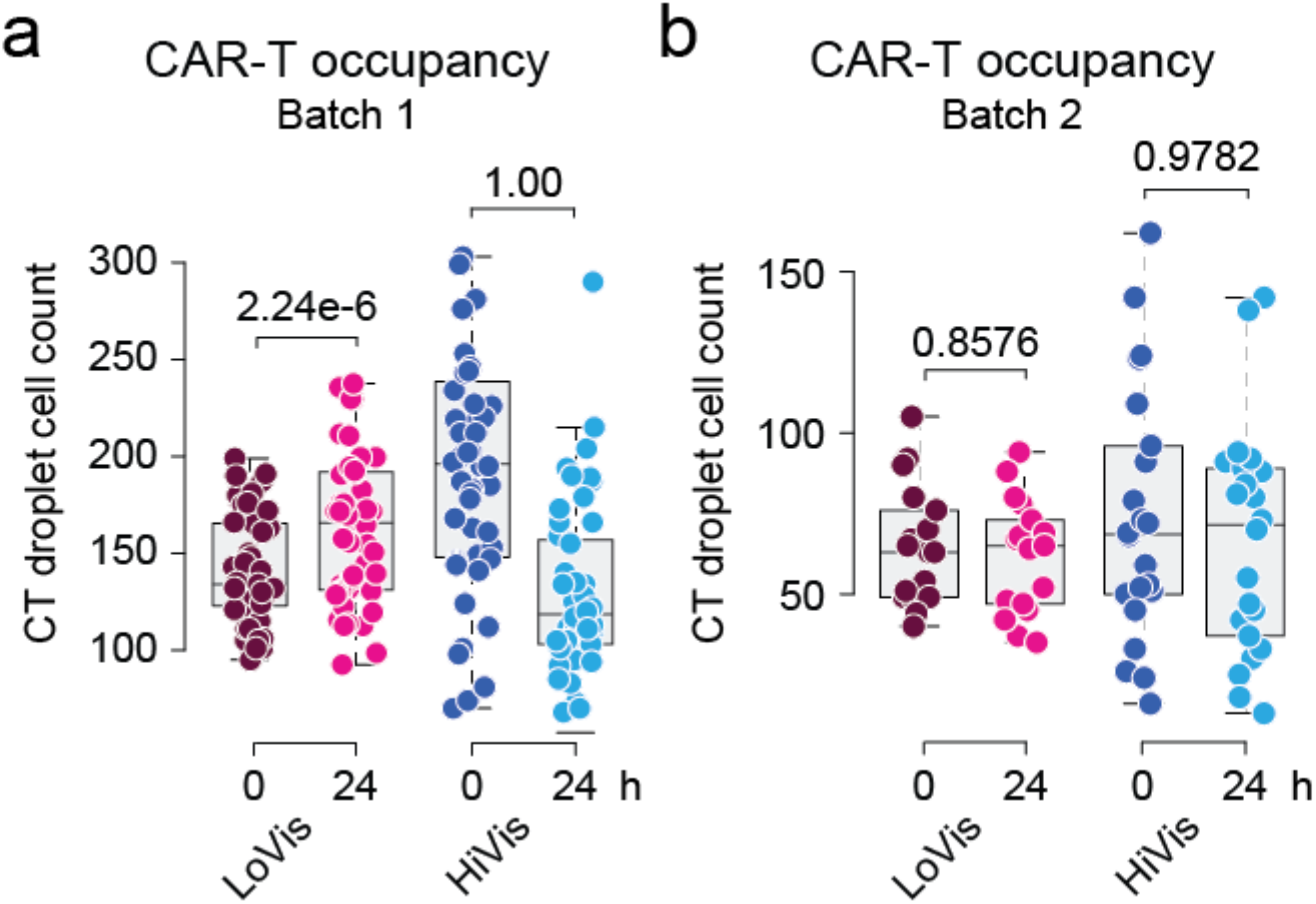
CAR-T cell expansion analysis. (**a**,**b**) Analysis of CAR-T occupancy before and after treatment on chip. Analysis of CAR-T occupancy before and after treatment on chip was performed. CAR-T droplet cell count was plotted against treatment time; each dot represented one well; the thick black line, grey box and dotted error bars of the box plot indicated median CAR-T droplet cell count, interquartile range between -1σ and 1σ, and range of all wells, respectively; the numbers indicated p-values of 1-tailed *t* test.

### Bioimaging study of CAR-T cells in action

Lastly, time-lapse imaging of CAR-T cells in action was conducted to dissect the paradigm of CAR-T cell killing in order to understand our observed mechanism of action. CAR-T and Nalm6 cells were mixed and loaded in a small hole in polydimethylsiloxane (PDMS) fabricated on a 35 mm Petri dish (Figure S3a) to reduce sample input and cope with microscope settings. Culture medium on top of the PDMS maintained the volume to avoid drying out from evaporation. Cell movement caused by natural heat convection in the 37°C humidified incubator and wetting between the unbound surfaces of PDMS and culture dish was amplified in the small volume, resulting in a mean flow rate of 1.4 μm/min (Figure S3b), which was much slower than naive T cells moving at an average speed of 10-15 μm/min on dish ^21^ or at a peak velocity of 40 μm/min in lymph nodes ^22^. Although the cell movement in our assay was much slower than the minimum blood flow of 90 cm/min in human fingers ^23^ or 60 μm/min in lymph vessels ^24^, this flow rate was sufficient to help us distinguish whether CAR-T cells were hunters or gatherers. Unlike cytotoxic T cells ^21,22^ and CAR-T cells ^25^ that moved on solid substrates, CAR-T cells suspended in fluid medium was carried by bulk flow and caught their prey using cellular protrusions that constantly emerged in active cells (Movies S1 & S2). Alternatively, active cell migration on dish by amoeboid locomotion was also observed (Movie S2), suggesting that CAR-T cells are predominantly gatherers in fluid, but also act as hunters on solid substrate. Regardless of the predation mode, the catches were swift and active cell-cell interaction continued along the bulk flow (Movies S1 & S2).

Image montages were extracted from the movies for detailed analysis of the process of CAR-T killing.

Scanning referred to the process of CAR-T cells searching for their targets. Active CAR-T cells constantly triggered cellular protrusions in all directions (Figure 7a), hinting of T cell resetting ^26^ and serial triggering ^27^. Bypassing cells were scanned randomly and rapidly, whereas the suspect was caught swiftly (Movies S1 & S2). The swift catch suggested of the involvement of high-affinity ligand-receptor interaction, which was assumed to involve the antigen-antibody pair of CD19-FMC63-scFv in our case, with a reported IC_50_ of 15.6 nM ^28^. Once a prey was caught, the CAR-T cell would attach to its prey via its cellular protrusion(s) (Figure 7a). If the prey was considered a non-target, the CAR-T cell would release it (Figure 7a), suggesting that proofreading also took place *in vitro*.

**Figure 7.**
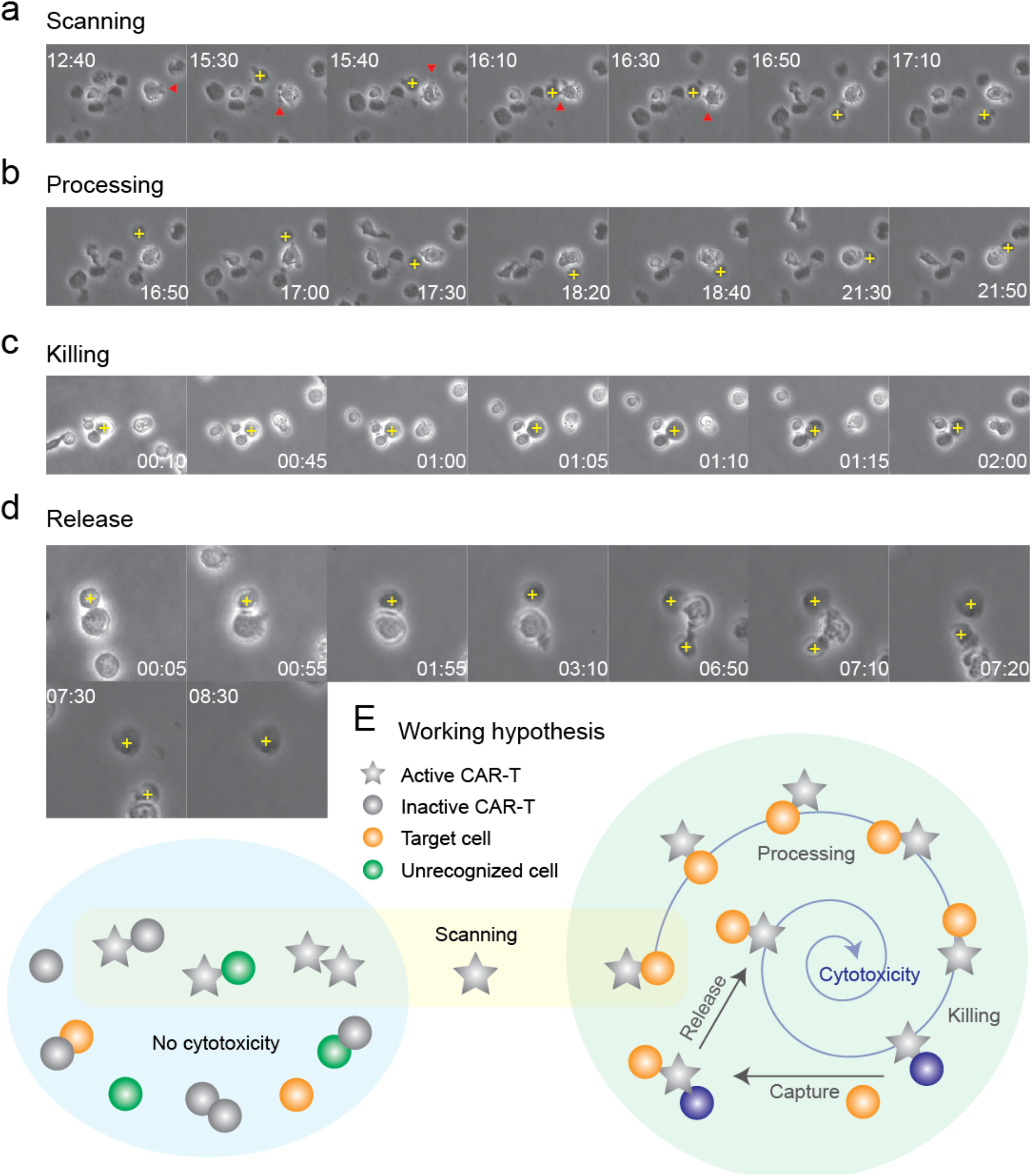
CAR-T killing paradigm. (**a-d**) Bioimaging study of CAR-T killing. CAR-T cells were mixed with Nalm6 cells for phase contrast imaging under 20x objective on Nikon Biostation. Sequential images of CAR-T cells undergoing the processes of scanning, processing, killing and release was composed as a montage by ImageJ. Yellow cross indicated target cell; red arrows indicated filopodia. (**e**) Working hypothesis of CAR-T killing. Active CAR-T cells scan for prey randomly, followed by processing, killing, capturing of a new prey and releasing the victim.

Processing of a prey to distinguish between target and non-target cell is critical for immune homeostasis. During processing, spinning of the prey around the CAR-T cell was observed (Figure 7b). However, this phenomenon could be interpreted in two ways: (a) attachment of the CAR-T cell to its prey induced a momentum that made the two cells spun together; or (b) the CAR-T cell remained stationary, and pulled its prey to rotate around itself by alternating surface adhesion (Figure S3c), which might involve cytoskeletal rearrangement ^29^. In scenario (a), spinning was simply a consequence of physical momentum and had no biological implication. On the contrary, scenario (b) suggested of surface receptor scanning during predator-prey interplay, which might involve autoimmunity prevention and/or co-stimulatory signal activation. A similar processing phenomenon was observed in Jurkat cells (data not shown) and in CAR-T cells *in vivo* ^25^, but not in other cell types that we investigated (data not shown). Hence, we are prone to believe that surface receptor scanning occurred, forming immunological synapses that developed into kinapses ^21^ to activate CAR-T cells. Nevertheless, identification of the underlying mechanism requires ultra-fast high-resolution imaging of actin dynamics, which had not been successful. Contrariwise, direct killing without spinning was also observed (Figure S3d). Nevertheless, processing with or without spinning consumed more than 1 h, and in both batches of CAR-T cells in this study, suggesting that the lengthy processing was not correlated to cytotoxic activity. In stark contrast, CAR-T cells took approximately 25 min to kill its target in bone marrow ^25^, whereas antigen presentation by B cell *in vitro* ^30^ or dendritic cell *in vivo* ^22^ consumed less than 1 h. Thus, it remained elusive if incorporation of three co-stimulatory receptors CD28, CD3ζ and TLR2 (Figure 1a) in our CAR-T cells extended the processing time.

After confirming the target, killing would take place where the prey would burst in necrosis-like cell death within 2 h (Figure 7c). However, the dead cell would not be released until a new prey was caught (Figure 7d). Nevertheless, cell release completed within 30 min (Figure 7d). The stringent adhesion of CAR-T cells to its victim until encountering a new prey and the short release time afterwards suggested that the turnover time of CAR-T cells relied on the frequency of predator-prey interaction, at least *in vitro*. This observation supports our quantitative on-chip analysis that CAR-T cytotoxicity dropped in large clusters and in high viscosity medium, which conferred to micro- and macro-scale cellular immobility, respectively. Additionally, our observation of active CAR-T killing in large clusters in low viscosity medium suggested that cell motility overcame micro-scale cellular immobility in extended cell aggregates. In support of our observation, CAR modification to reduce off-rates was shown to increase cytotoxicity ^31^.

To summarize, the procedures of CAR-T cytotoxicity involved four steps: (1) scanning of prey to identify its target, followed by release of prey if mistaken; (2) processing of prey to activate cytotoxic program of CAR-T cells; (3) killing the prey by necrosis-like cell death; and (4) release of the dead cell after a new prey was caught (Figure 7e).

## Discussion

In this study, we introduced a novel analytic method to assess CAR-T cytotoxicity *in vitro*. However, we have not accomplished cell type identification to enable cytotoxicity test on primary cancer cells, unless CAR-T cells are clinically approved to contain a fluorescence label. Nevertheless, our assay provides a new tool to study cellular interaction between cancer cells and CAR-T cells *in vitro*.

Furthermore, we made significant improvements to our previous assay ^7^. Specifically, we replaced the parameters of mean droplet cell viability and range of droplet cell viability with two ratiometric parameters, namely the cell ratio and well ratio, respectively. The virtue of using ratiometric parameters is to eliminate the impact of different sample size across replicate wells to robustly evaluate CAR-T cytotoxicity as a population and in a population. This also unifies the scale to dissect the mechanism of action among different phenotypes, e.g. CAR-T cytotoxicity in different cell categories. Phenotype analysis provides unique insight for biological studies. Mechanistically, our results showed that high fluid viscosity and formation of larger clusters impedes CAR-T cytotoxicity, which confer to cellular immobility on the macro- and micro-scale respectively. However, given the fact that CAR-T cells were still actively killing Nalm6 cells in large clusters in low viscosity medium, we deduced that cell motility could overcome micro-scale cellular immobility. Notably, Nalm6 cluster shuffling between categories was anticipated but could not be quantified under current circumstances. For instance, the Nalm6 cells increased in large clusters at 24 h post-treatment in low viscosity medium in Batch 2 CAR-T cells (Figure 5b) probably came from other cell categories because cells could not appear from none. However, the overall unchanged distribution of Nalm6 occupancy among categories at 24 h post-treatment ruled out massive, unidirectional cluster shuffling, if any had occurred.

On the other hand, Nalm6 occupancy among different cell categories was more evenly distributed in Batch 1 CAR-T cells (Figure 3b) as compared to Batch 2 CAR-T cells (Figure 5b) in low viscosity medium upon chip loading. This observation raised the discrepancy that active CAR-T cells are prone to aggregates whereas large clusters impede CAR-T cytotoxicity. To explain this phenomenon, the possibility of cell aggregation in ambience due to experimental timing was first ruled out. Next, we sought to solve this issue by studying CAR-T cells in action. Time-lapse imaging showed that active cells rapidly scanned prey (Movie S1), which confers to higher probability of cell cluster formation in the more active Batch 1 CAR-T cells. Furthermore, the rigorous catch-and-release of prey by CAR-T cells posits a higher chance of cluster formation in the more active Batch 1 CAR-T cells, provided that the frequency of target cell encounters is conserved between batches due to similar loading cell densities.

Another interesting observation in this study is the dilemma between CAR-T cytotoxicity and CT:N6 ratio on plate and on chip. Although luciferase assay depicted positive correlation between CT:N6 ratio and CAR-T cytotoxicity on plate (Figure S1), on-chip screening showed negative correlation in Batch 1 (R = 0.3541, p = 0.0136) and Batch 2 (R = 0.2435, p = 0.0441) CAR-T cells in low viscosity medium (Figure S4a & S4c). No correlation between CT:N6 ratio and CAR-T cytotoxicity was observed in high viscosity medium in Batch 1 (R = -0.1250, p = 0.3973) and Batch 2 (R = 0.0392, p = 0.3374) CAR-T cells in high viscosity medium (Figure S4b & S4d), in concordance to the observed poor cytotoxic activity under this fluid condition. Initially, we postulated that exponential decay of bioluminescence exaggerated the number of dying CAR-T cells on plate. After that, comparison between label-free analysis and analysis considering EH1-labelled cells in the same dataset showed that absolute cell number counting was less susceptible to signal fluctuation as compared to labelled cells. Lastly, our observation of intensive bypassing cell scanning by CAR-T cells as gatherers suggested that increasing CT:N6 ratio in a confined volume of the droplet will reduce the probability of Nalm6 cell encounters. Taken together, we hypothesized that these three factors constituted to the discrepancy of the correlation between CT:N6 ratio and CAR-T cytotoxicity using different assessment methods. Nonetheless, the bulk droplet CT:N6 ratio becomes less meaningful on chip when the mechanism of action could be evaluated via phenotype analysis.

Lastly, label-free cytotoxicity analysis is achieved in this assay by acquiring images before and after treatment. Additionally, dual time point acquisition accomplished discrimination of CAR-T cytotoxicity into two processes: (1) CAR-T killing of cancer cells; and (2) CAR-T cell expansion. This feature will be enhanced by the virtue of defined solvent composition in *in vitro* assays, that is, to dissect CAR-T cytotoxic activity by varying cytokines and chemokines, *etc*. Moreover, generalization of our two-cell type model into multi-cell type model by diversification of fluorescence labeling of each cell type will facilitate the study of complex cell-cell interactions at ease. In addition, the frequency of data acquisition can be increased up to every 4 h under current circumstances, suggesting that phenotype-based outcomes other than cell death can be assessed using this method. Choice of appropriate reporter(s) not only enables phenomenal observations, but also provides mechanistic insight on the biological pathways under investigation. Nevertheless, the current assay is technically restricted to genetically engineered cells, unless novel cell labeling methods or cellular monitoring methods are applied. Another drawback of this method is the restriction to cells that are adaptable to suspension culture. A better understanding of the physical properties of tumor and biogels will be needed to establish *in vitro* assays for adherent cells ^16,17^.

## Conclusion

We presented a novel droplet microfluidic assay for analyzing CAR-T cytotoxicity on the aspects of cancer cell reduction and CAR-T cell expansion, and provided insight on the mechanism of action via phenotype analysis. This method provided unprecedented opportunity to understand cell-cell interaction quantitatively on the single cell level. Consequently, this method can be generalized to apply to any cell-kill-cell experiment provided that all cells are empirically distinguishable.

## Acknowledgements

Acknowledgements to University of Macau Development Foundation for financial support on our patent application. This work was supported by The Science and Technology Development Fund, Macau S.A.R. (File no. 137/2016/A) granted to A.H.W., and The Science and Technology Development Fund, Macau

S.A.R. (File no. 110/2016/A3) granted to Y.J., P.I.M. and A.H.W.; the Strategic Priority Research Program of the Chinese Academy of Sciences Grant No. XDB19030205, Guangdong Provincial Applied Science and Technology Research & Development Program No. 2016B020237006, Guangzhou Science and Technology Plan Project No. 201907010042 and 201904010473, and Frontier Research Program of Guangzhou Regenerative Medicine and Health Guangdong Laboratory Grant No. 2018GZR110105003 granted to P.L.; the UM Chair Professor Fund No. CPG2020-00004-FHS granted to C.D.. Infrastructure for microfluidic chip fabrication and cell culture was provided by Y.J. and C.D., respectively. J.K.W. was partially financially supported by C.D..

## Author Contributions

This project was conceived by A.H.W. through collaboration with P.L.. A.H.W. designed the study and supervised J.K.W. and H.W. on experiments and bioinformatics analysis, respectively; P.L. supervised J.S.. Preparation and FACS characterization of CAR-T cells were performed by J.S.; the luciferase and chip assays were performed by J.K.W.; the bioinformatics analyses and data visualization were performed by H.W.; the bioimaging study was performed by A.H.W.; statistical analysis was performed by H.W. and A.H.W.; Y.J. provided the chip design; L.J. shared perspectives on the findings. All authors prepared and approved this manuscript.

## Data and Materials Availability

All data is available upon request from the corresponding authors.

## Supplementary Figures

**Figure S1.**
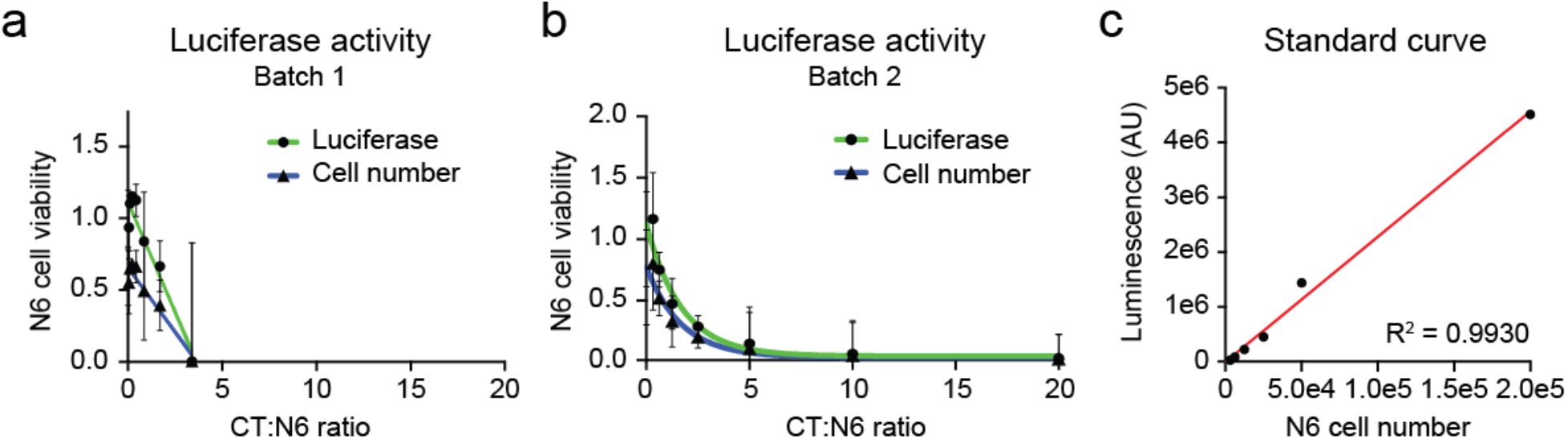
CAR-T cell characterization. (**a-b**) Luciferase assay of Batch 1 and Batch 2 CAR-T cells. Nalm6 cell viability was plotted against CT:N6 ratio; each dot represented mean Nalm6 cell viability of three replicate experiments; the lines indicated non-linear regression of the data points by GraphPad Prism; the error bars denoted standard deviation of the mean. (**c**) Standard curve of luminescence against Nalm6 cell numbers. In order to compare results between conventional luciferase assay and our chip assay, we performed serial dilution of Nalm6 cells to plot the standard curve. Luminescence was plotted against Nalm6 cell number; each dot represented the mean of three replicates; the red line indicated linear regression of the data points by GraphPad Prism.

**Figure S2.**
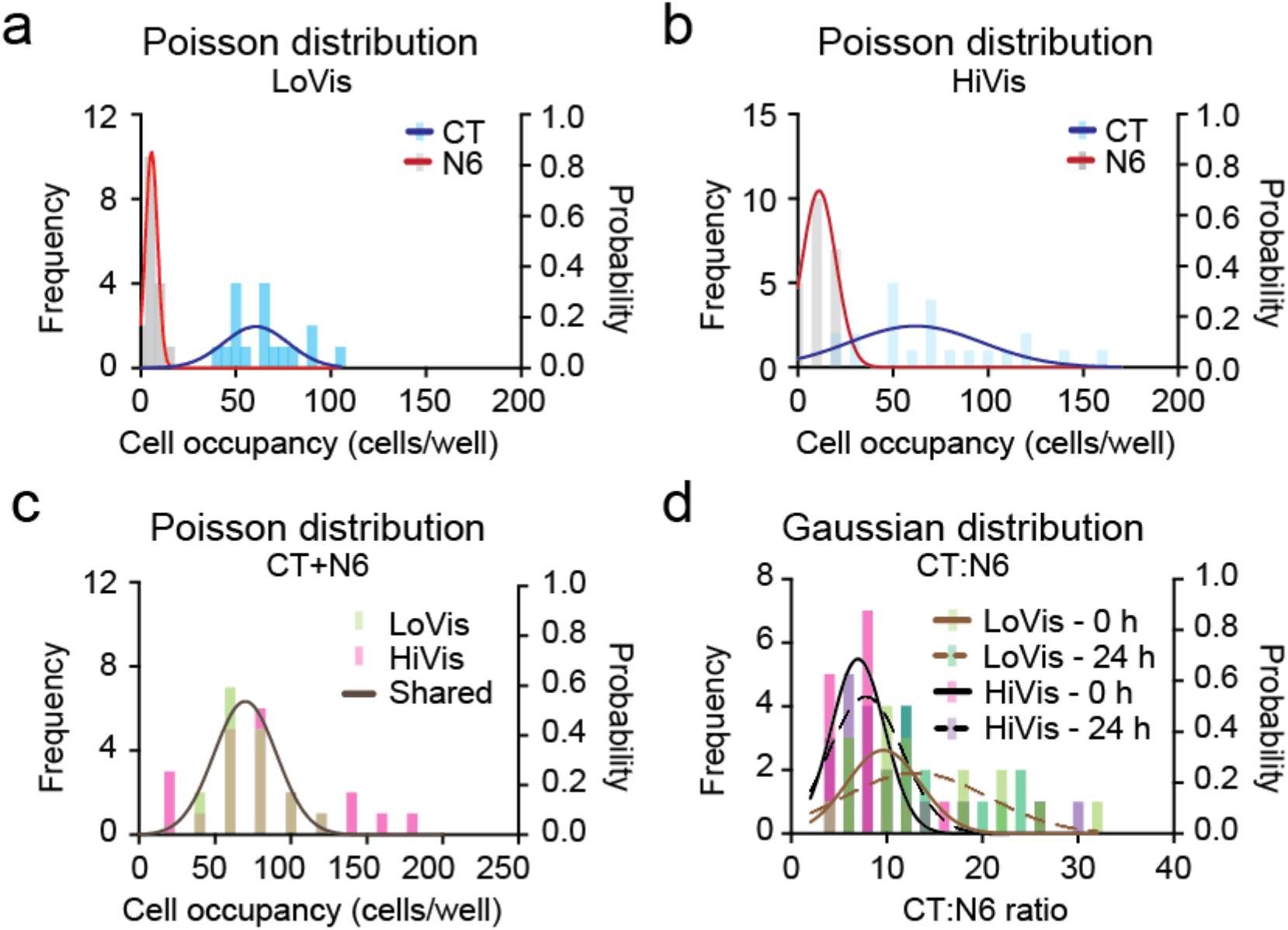
Cell occupancy analysis of Batch 2 CAR-T cells. (**a-c**) Poisson distribution of Batch 2 CAR-T and Nalm6 cells on chip. Frequency distribution analysis of CAR-T (CT) and Nalm6 (N6) cells in low viscosity medium (LoVis) and high viscosity medium (HiVis) on chip at 0 h was performed. Distribution frequency and probability was plotted against cell occupancy in all wells on chip; each column represented one data grid automatically classified by GraphPad Prism; the lines indicated Poisson distribution curves generated by GraphPad Prism. (**d**) Gaussian distribution of CT:N6 ratio on chip. Frequency distribution analysis of CT:N6 ratio under both fluid conditions before and after treatment was performed. Distribution frequency and probability was plotted against CT:N6 ratio in all wells on chip; each column represented one data grid automatically classified by GraphPad Prism; the lines indicated Gaussian distribution curves generated by GraphPad Prism.

**Figure S3.**
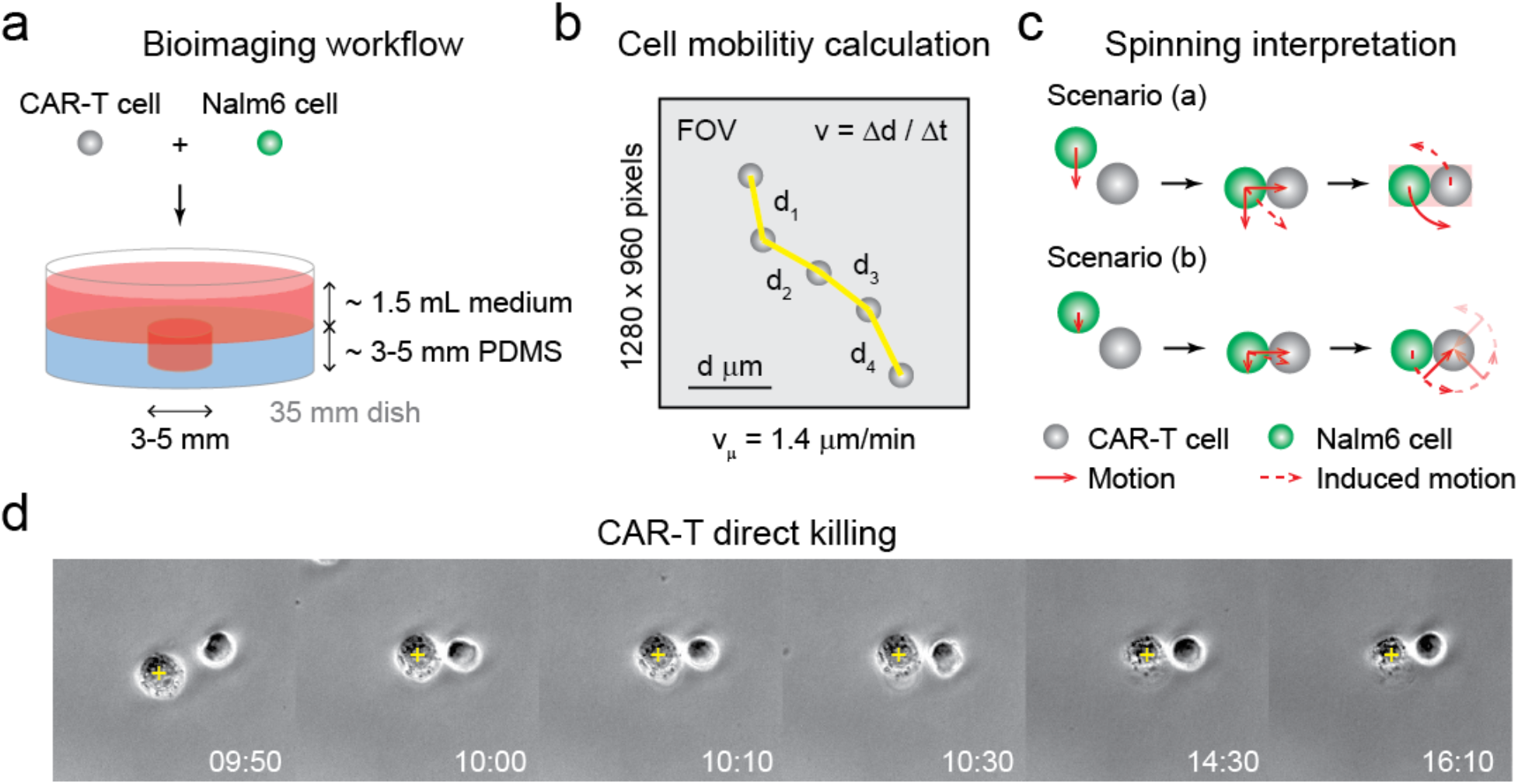
Bioimaging study of CAR-T killing. (**a**) Bioimaging workflow and setup. CAR-T and Nalm6 cells were loaded into the PDMS well fabricated on a 35 mm cell culture dish, followed by time-lapse imaging on Nikon Biostation at 5-10 min intervals under 20× phase contrast. (**b**) Cell mobility calculation. Cell mobility was calculated by mean displacement divided by time in 3 cells from 3 different regions of interest (ROIs). (**c**) Schematic interpretation of cell spinning during processing. Scenario (a) hypothesized that spinning was a result of momentum during cell adhesion, whereas Scenario (b) hypothesized that spinning was a result of alternating surface adhesion, triggered by surface receptor scanning. (**d**) Evidence of CAR-T direct killing. CAR-T cells were mixed with Nalm6 cells for phase contrast imaging under 20× objective on Nikon Biostation. Sequential images of CAR-T cells killing its prey without spinning was composed as a montage by ImageJ. Yellow cross indicated target cell; time was indicated in hh:mm format.

**Figure S4.**
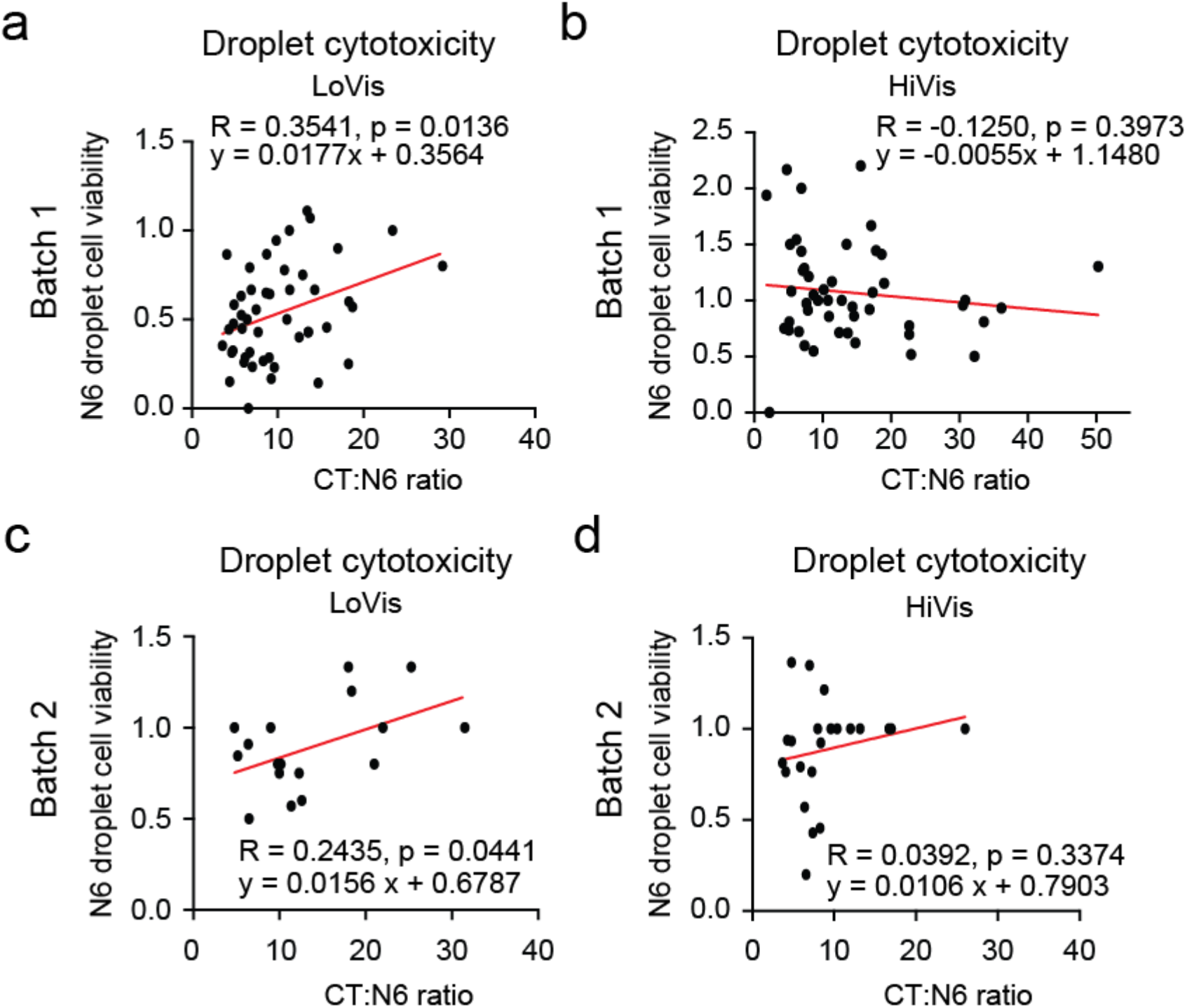
Correlation analysis of CAR-T cytotoxicity and CT:N6 ratio under different fluid conditions. (**a-d**) Correlation analysis of CAR-T cytotoxicity and CT:N6 ratio of Batch 1 and Batch 2 CAR-T cells under different fluid conditions. Nalm6 droplet cell viability was plotted against CT:N6 ratio; each dot represented one well and the red line indicated linear regression of the data points by GraphPad Prism.

## Supplementary Movies

**Movie S1**. Time-lapse imaging of Batch 2 CAR-T cells and Nalm6 cancer cells.

**Movie S2**. Time-lapse imaging of Batch 1 CAR-T cells and Nalm6 cancer cells.

## Supplementary Data

**Data S1**. Supplementary Tables S1-S7.

**Table S1**. One-way ANOVA analysis of Nalm6 droplet cell count against different cell categories at 0 h in Batch 1 CAR-T cells.

**Table S2**. Two-way ANOVA analysis of Nalm6 droplet cell count in different cell categories against well position at 0 h in Batch 1 CAR-T cells.

**Table S3**. Two-way ANOVA analysis of Nalm6 droplet cell count difference between 0 h and 24 h in different cell categories against well position in Batch 1 CAR-T cells.

**Table S4**. Pearson correlation analysis of Nalm6 droplet cell viability between analyses with and without consideration of EH1 labelling in Batch 1 CAR-T cells.

**Table S5**. Pearson correlation analysis of Nalm6 droplet cell count between analyses with and without consideration of EH1 labelling at 0 h and 24 h in Batch 1 CAR-T cells.

**Table S6**. Comparison of CAR-T expansion between low viscosity medium and high viscosity medium in Batch 1 CAR-T cells.

**Table S7**. Comparison of CAR-T expansion between low viscosity medium and high viscosity medium in Batch 2 CAR-T cells.

